# Geographic distribution of genetic diversity of *Heterocephalus glaber* analyzed using whole genome sequencing and a chromosome-scale genome assembly

**DOI:** 10.1101/2025.07.22.666215

**Authors:** Kevin M. Wright, Pranidhi Sood, Elena D. Zemlemerova, Danila S. Kostin, Nicole L. Fong, Nelda Yi, Andrea T. Ireland, Irene Lam, Kaitlyn N. Hardell-Lewis, Megan Smith, Jackie Villalta, Calvin H. Jan, Margaret A. Roy, David Botstein, Leonid A. Lavrenchenko, J. Graham Ruby, Rochelle Buffenstein

**Affiliations:** Calico Life Sciences LLC, 1170 Veterans Blvd, South San Francisco, CA 94080, USA; Actio Biosciences 11202 El Camino Real, Suite 100. San Diego, CA, 92130; Department of Mammalian Microevolution, Severtsov Institute of Ecology and Evolution, Russian Academy of Sciences, Moscow 119071, Russia; Department of Biological Sciences, University of Illinois, Chicago 845 W Taylor, Chicago, IL 60607, USA

## Abstract

Naked mole-rats (*Heterocephalus glaber*) are a species of rodent endemic to the Horn of Africa, notable among mammals for their long lifespans, resistances to a variety of stresses, and eusocial mating behavior. Though their natural range extends across large portions of Kenya, Ethiopia, Somalia, and Djibouti, the large majority of genetic and genomic analyses focus on Kenyan specimens. Here, we constructed a chromosome-scale reference genome assembly for *H.glaber*, along with new reference assemblies of both the Damaraland mole-rat (*Fukomys damarensis*) and guinea pig (*Cavia porcellus*) genomes to aid annotation. We leveraged our *H.glaber* assembly, along with modern whole-genome sequencing, to characterize the genetic diversity of specimens deriving from Kenya, southern Ethiopia, and eastern Ethiopia. We found the Kenyan and southern Ethiopian specimens to be closely related to each other and highly diverged from eastern Ethiopian specimens. We also found specimens collected from nearby locations in southern Ethiopia to be more closely related to Kenyan specimens than to each other. This unexpected distribution of shared genetic diversity highlights the importance of local migration barriers to gene flow in wild *H.glaber* populations.

## Introduction

Naked mole-rats (Rodentia; *Heterocephalus glaber)* are renowned for several unique behavioral, physiological and metabolic traits (Buffenstein, Park and Holmes 2021-book). These eusocial, subterranean-dwelling rodents exhibit extreme longevity (Ruby et al., 2024; Buffenstein 2005) with a concomitant negligible senescence phenotype Buffenstein 2008; Can et al., 2022), and fail to exhibit demographic aging, maintaining a constant mortality hazard for more than three decades (Ruby et al, 2018, Ruby et al., 2024). They show pronounced tolerance of hypoxia and hypercapnia (Park et al., 2017; Zion et al) and also exhibit exceptionally low cancer incidence relative to that shown by most other mammals (Buffenstein 2008; Delaney et al. 2016; Hadi et al, 2021). Their immune system paradoxically lacks natural killer cells -that in other species play a crucial role in the defense against cancer (Hilton et al., 2019) - but unusually has a substantial circulating population of cytotoxic gamma delta T (γδ T) cells (Lin et al., 2024). The molecular mechanisms behind these unusual features remain, however, poorly understood.

Naked mole-rats exhibit extreme variation in lifetime reproductive success as well as cooperative care of siblings, features that has been described as a eusocial breeding system (Jarvis, 1981; Holmes & Goldman, 2021, Buffenstein et al., 2022), with fewer than 1% of individuals reproducing over their lifespans (Jarvis, 1991). Indeed, most naked mole-rat colonies only have one breeding female, who will birth all of the offspring in the colony (Jarvis, 1981), and a handful of breeding males (Faulkes *et al*., 1997). These rodents naturally live in isolated underground colonies, characterized by low rates of immigration and emigration (Jarvis, 1985). New colonies are thought to primarily arise from the splitting up of existing colonies and subsequent matings among closely related individuals. Naked mole-rats are described as facultative inbreeders because the breeding female will mate with the founder male and, as the colony grows, additionally may mate with a few of her male offspring (Faulkes and Bennett 2021). These features are thought to have resulted in high-within colony relatedness (Reeve et al.,1990; Honeycutt et al., 1991; Ingram et al., 2015).

Simultaneous appearance of fossils resembling present-day *Heterocephalus*, together with extinct bathyergid ancestors in Miocene deposits, supports an ancient origin for this genus, no later than 18 million years ago. This fossil evidence, together with molecular clocks, suggest it to have been the earliest phylogenic branch to diverge within the African mole-rat lineage (Faulkes and Bennett, 2021). Phylogeographic analyses suggest their distribution to have been predominantly influenced by landscape evolution involving the formation of physical barriers (Hess et al, 2022), as well as ecological and climatic modifications associated with the formation of Africa Rift Valley. The natural species range of naked mole-rats spans the Horn of Africa, extending from the southern border of Kenya, across the Somali-Masai biome, spanning Ethiopia and reaching as far north as Djibouti (Zemlemerova et al, 2020; Bennett and Faulkes 2000; Faulkes and Bennett 2021). Most field studies and sampling of naked mole-rats have occurred in Kenya (Jarvis 1981; Brett 1991a,b; Jarvis and Sale 1971). The first genetic study of this species, using specimens from southern Kenya, measured low levels of genetic variation and concluded their eusocial mating system was responsible for a high inbreeding rate (Reeve et al. 1990). However, a follow up study with broader geographic sampling found greater genetic diversity in northern Kenya (Ingram et al. 2015), suggesting migration-induced population bottlenecks at the edge of the species range as responsible for the low genetic variation observed in southern Kenya. For this species, the use of genetics to inform population history and demography has heretofore been limited to analysis of a restricted set of autosomal and mitochondrial markers (Zemlemerova et al, 2020; 2021).

Here, we examined the natural geographic distribution of genetic diversity amongst naked mole-rats using polymorphisms that were densely annotated from whole-genome sequencing, using animal specimens that originated from Kenya as well as southern and eastern Ethiopia. We found the Kenyan *H.glaber* to be closely related to southern Ethiopian specimens and highly divergent from eastern Ethiopian populations. Within southern Ethiopia, we found geographically proximal populations to be highly diverged, and to be more closely related to the more distant Kenyan populations than they were to each other.

In order to facilitate our genetic analyses, a new, chromosome-scale genome assembly for *H.glaber* was constructed, by using a combination of high-coverage short-read, long-read, and optical-mapping data. This new resource, along with gene annotations and underlying data, are described and provided along with this manuscript. To additionally support those annotation efforts, we generated and provide new reference assemblies of both the Damaraland mole-rat (*Fukomys damarensis*) and guinea pig (*Cavia porcellus*) genomes.

## Results

### *H.glaber* reference genome assembly and gene annotation

Our workflow for genome assembly and annotation is described below and schematized in Supplemental Figure 1, and assembly progress across each step of the process is reported in Supplemental Table 1. Multiple data modalities were leveraged - including short-read sequencing, long-read sequencing, chromosome-seq and optical mapping - to construct a chromosome-length genome assembly of the naked mole-rat, following the Vertebrate Genome Project assembly protocols (Rhie et al. 2021). Primary assembly and polishing were done with PacBio long-reads and BioNano optical reads. Secondary scaffolding and polishing were done with the 10X linked-reads sequencing in which long DNA molecules are prepared in a bar-coded library and sequenced using a highly accurate short-read sequencer. Deviations from the standard VGP1.6 pipeline are described in the Methods. Finally, scaffolds likely to derive from the same chromosome were grouped using sequence from flow-sorted chromosomes (Soifer et al. 2020).

The diploid genome of *H.glaber* includes 29 pairs of autosomes plus sex chromosomes (Deuve et al, 2006) and is 5.8 picograms of DNA (Gallardo et al, 2003), which corresponds to ∼2.84 gigabases (Gb) in a haploid genome (Dolezel et al, 2003). Our assembly included 2.53 Gb of sequence (2.56 Gb including scaffold-linking N’s) and therefore likely captured the majority of the genome. The 408 contigs were joined into 130 scaffolds, with N50 lengths of 22.88 megabases (Mb) and 68.60 Mb (69.35 Mb including scaffold-linking N’s), respectively (Figure 1A). A theoretical distribution of chromosome lengths based on the measured genome size and karyotype (see Methods) had an N50 of 102.3 Mb (Figure 1A). The less than two-fold difference between scaffold and karyotype-based N50 values suggested that scaffolds often traversed entire chromosome arms This was validated by clustering 58 of the largest scaffolds into 30 groups using sequence data from flow-sorted chromosomes (Figure 1B; Supplemental Table 2). The scaffold groups had an N50 of 97.30 Mb, close to the theoretical N50 for entire chromosomes (Figure 1A).

**Figure 1.**
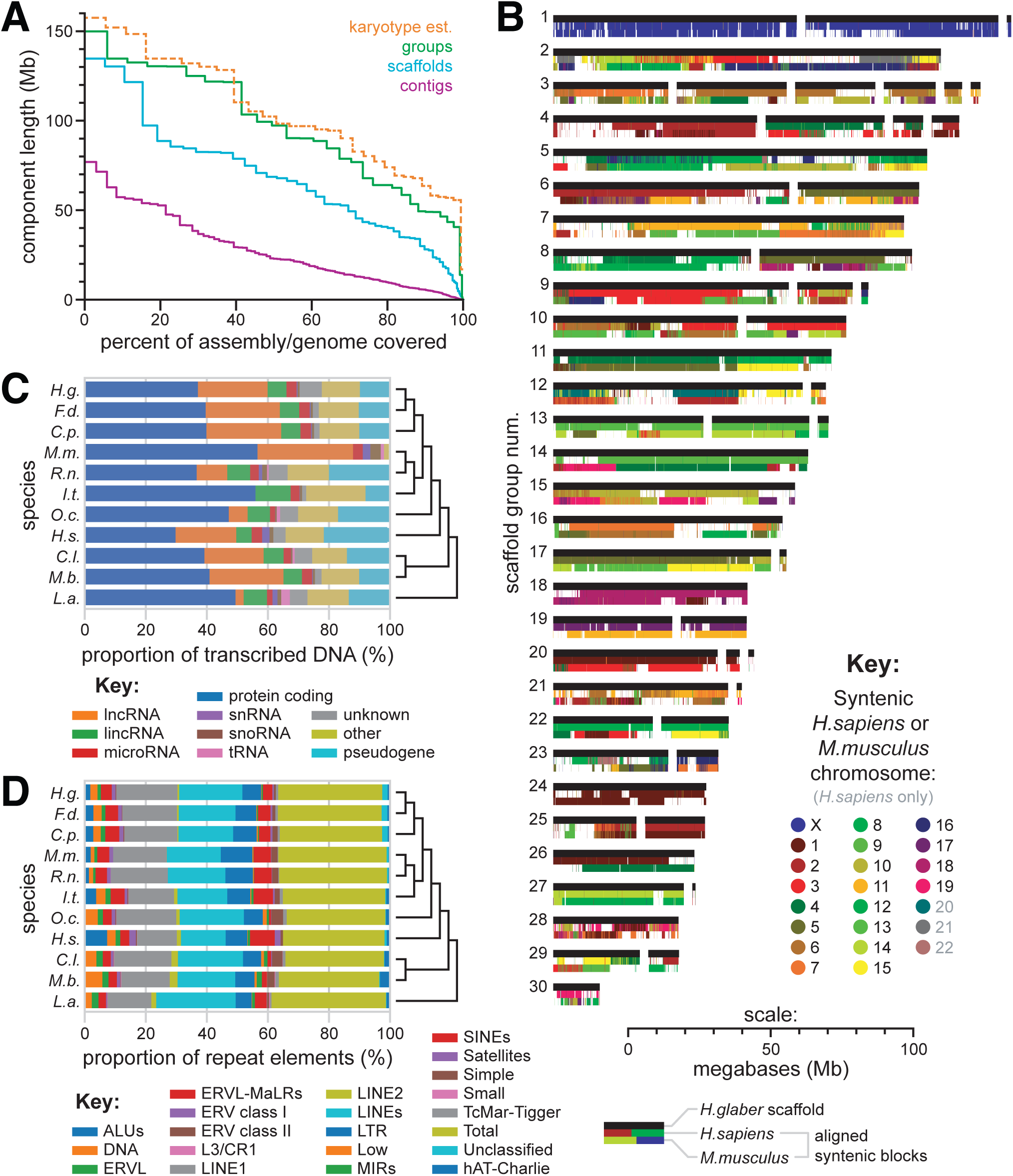
A. Relative length distributions for contigs (magenta), scaffolds (cyan), and scaffold groups (green) in our reference *H.glaber* genome assembly. In orange: the relative length distributions for chromosomes, based on analysis of karyotypes. B. Scaffold groups, based on sorted chromosome-seq. Black bars represent scaffolds, drawn to scale with their lengths. Colored bands below each scaffold bar indicate regions of syntenic alignment to human (middle band) or mouse (bottom band) reference genomes, color-coded by the aligned human/mouse chromosome. C. Proportion of transcribed DNA dedicated to various gene/transcript categories, as annotated by CAT, across all species included in our multi-genome alignment. D. Distribution across classes of repetitive elements labelled by RepeatMasker, as annotated by CAT, across all species included in our multi-genome alignment.

We annotated the *H.glaber* genome using the Comparative Annotation Toolkit (CAT) (Fiddes et al. 2018), run with the highly manually curated genomes of mouse and human as well as *ab initio* searches for genes specific to the less well annotated lineages: rat, squirrel, elephant, rabbit, bat and dog (Supplemental Figure 1). Genomes for the Damaraland mole-rat *Fukomys damarensis* and guinea pig *Cavia porcellus* were also included. For the latter two, due to their taxonomic proximity to *H.glaber*, we generated our own genome assemblies using 10X linked-read data (see Methods).

The CAT annotation pipeline performed whole genome alignments in Cactus (statistics provided in Supp. Table 3; whole-genome alignments provided as Supp. Data Files 3) and predicted gene family identities using comparisons to the well-annotated mouse and human genomes. Gene annotations from the mouse and human genomes were projected onto all other species. Those conservation-based predictions were supplemented and validated using *H.glaber* gene expression data from both Illumina short reads and PacBio long reads. Repetitive elements were similarly annotated across all ten genomes (see Methods for details). The proportions of both gene and repeat element classes were similar across the ten mammalian genomes included in the CAT analysis (Figure 1C,D).

That assembly exhibited extensive synteny with the human and mouse genomes (Figure 1B) and achieved a complete BUSCO score of 93.2% (Simão et al, 2015), suggesting its structure to be largely correct. Using our gene annotations and whole-genome alignments, we mapped syntenic blocks to human and mouse chromosomes onto our genome assembly (see Methods). Most scaffolds and groups were linked to individual orthologous chromosomes across large, tens-of-megabase blocks (Figure 1B). Notably, scaffold group 1 exhibited nearly exclusive synteny with the X chromosomes of both human and mouse, implying it to be the X chromosome of *H.glaber*.

### Polymorphisms discovered through whole-genome sequencing reveal kinship patterns within a captive *H.glaber* population derived from Kenya

Our captive *H.glaber* population descended from approximately 200 animals collected from six sites in eastern and southern Kenya in 1979 (Jarvis, 1985). Wild-caught animals from each collection site were maintained in captivity and interbred for multiple generations in order to maintain as much of the original genetic diversity as possible. Given that history, we sought to describe and quantify the genetic diversity that has been maintained from those founders, in order to gain insight into the amount of genome-wide genetic variance that exists across Kenyan haplotypes. To that end, we performed whole-genome sequencing on a large number of animals, selected to enrich individuals with distinct or unknown lineages.

The pedigree in Supplemental Figure 2 depicts the recorded genealogy of those captive animals. Given the eusocial structure of *H.glaber* society, nodes in this pedigree represent family-colonies (composed of a breeding pair and their offspring) and arrows represent the migration of an animal from its birth colony to become a breeder in a new family-colony. The records used to construct this pedigree were largely restricted to the previous 25 years, and in only a few instances was there enough information to trace relationships back to wild-caught progenitors.

In order to examine genetic relatedness between animals in this pedigree, we sequenced their genomes with short-reads to a median coverage of 20 to 26X (Supplemental Table 5). Figure 2A shows the subset of the pedigree linking those colonies whose members were sequenced. We identified SNP variants using the GenotypeGVCFs tool in GATK v4.0 (Poplin et al. 2017). High-confidence SNPs met the following criteria: bi-allelic variant, quality score greater than 30, sample missingness was less than 20%, minimum depth of coverage exceeded whichever of the two values was greater - the 5% genome-wide quantile or 6 reads, maximum depth was less than the 95% genome-wide quantile. Following this procedure, we identified between 2.8-4.1 million high-confidence SNPs per animal (Supplemental Table 5). Across all these animals, >8.1 million non-redundant polymorphisms were identified (Supp. Data Files 2).

**Figure 2.**
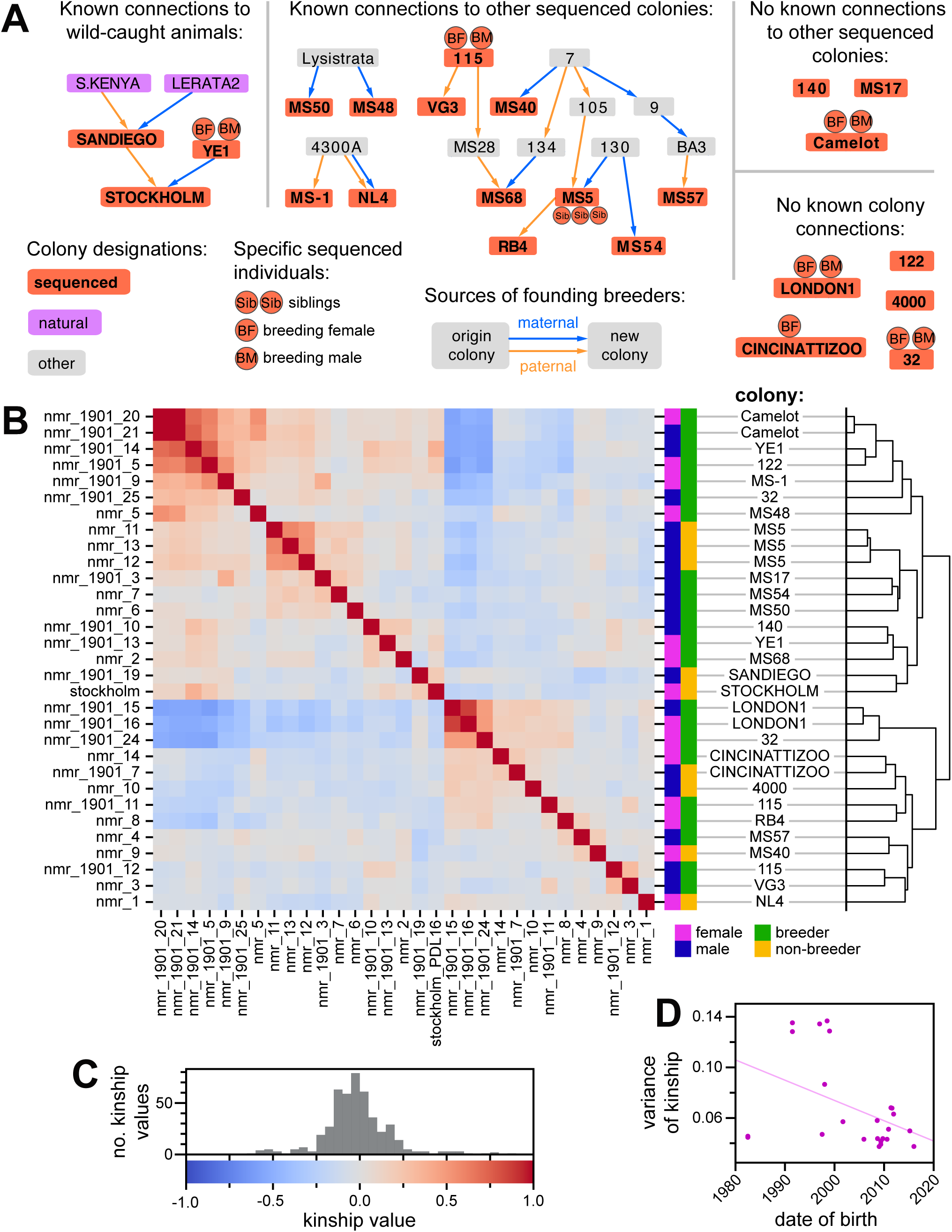
A. Partial pedigree of the Kenya-derived *H.glaber* collection. Nodes in this pedigree represent family-colonies (composed of a breeding pair and their offspring). Red nodes depict colonies that contain animals selected for whole-genome sequencing; abutting circles indicate when breeders or siblings were sequenced. Arrows show the origin-colony for the breeding male (orange) and female (blue) and the new-colony created by the union of these breeders. This panel includes networks linking sequenced colonies; the full pedigree is provided as Supplemental Figure 2. B. A heatmap of pairwise kinship values (*r^w^*) between the 31 sequenced *H.glaber* individuals from the Kenya-derived collection. Hierarchical clustering of animals based on kinship is shown on the right. The key for color intensity values for the heatmap is shown in panel (C). C. The key for color intensity values for the heatmap in panel (B), with a histogram showing the distribution of observed values. D. Plot of, for each animal, its date of birth (x-axis) versus the variance (y-axis) of its distribution of non-self kinship values (either rows or columns of panel B). Line shows linear regression (slope = −0.0016; y-intercept at 1980 = 0.106; slope SE = 0.00072; p-value = 0.036)

Whole genome sequencing data were used to estimate kinship (VanRaden 2008; Goudet, Kay, and Weir 2018) between all samples (Figure 2B). The majority of samples were from colonies without known pedigree connections, and estimates of kinship ranged from −0.2 to 0.2 (Figure 2C). In contrast, pairwise kinship values for the three full-siblings from colony *MS5* ranged from 0.521 to 0.587, similar to expected pairwise kinship of 0.5.

Additional pairs or clusters of animals without known pedigree connections nonetheless shared kinship exceeding expectations for full siblings (e.g. nmr_1901_14, from colony YE1, and nmr_1901_5, from unlinked colony 122, whose pairwise kinship was 0.75; Figure 2B; Supplemental Table 6). Many of these also shared lower-than expected kinship values across the sequenced individuals, including negative kinship values, which can indicate population structure (Figure 2B; see Methods for discussion of the meaning of negative kinship values). This population descends from animals collected six sites in Kenya (Jarvis 1985), found across regions known to be genetically distinct (Ingram et al. 2015). Although animals were cross-bred through the history of the collection, it can take multiple generations for linkage disequilibrium to decay, even between unlinked loci (Pfaff et al, 2001). We hypothesized that animals born earlier in the collection’s history possessed less-mixed genotypes, resulting in more extreme kinship values versus one another, in both the positive and negative directions. To test this hypothesis, the variance of kinship values for each was regressed against that individual’s date of birth (Figure 2D). The relationship was significantly negative (p-value = 0.036), with substantial admixture having been achieved after the year 2000.

Subsequent analyses were restricted to 24 animals by excluding all but a single animal from each such cluster sharing kinship above 0.5 (see Methods for details). The remaining animals carried >7.6 million (>94%) of the non-redundant polymorphisms discovered across the Kenya-derived cohort (Supp. Data Files 2).

### Polymorphism discovery in Ethiopian *H.glaber* samples reveals high genetic divergence for eastern Kenyan populations

We next sought to compare levels of genetic diversity within the Kenya derived population to four and six wild-caught animals from southern and eastern Ethiopia, respectively (Figure 3A; Supplemental Table 4; Zemlemerova et al., 2021). The genomes of the Ethiopian animals were short-read sequenced to a median coverage of 13-22X (Supplemental Table 5), and high-confidence polymorphisms were called using the same methods used for the Kenyan population. We identified 3.5 - 4.9 million sites with non-reference alleles per animal in the southern Ethiopian lines and 18.4 - 18.6 million sites per animal with non-reference alleles in the eastern Ethiopian lines (Supplemental Table 5). The sequencing of animals from southern and eastern Ethiopia added >3.9 million and >17.6 million non-redunant polymorphisms, respectively, to our discovery set, bringing the total (including Kenya-derived) to >30.1 million (Supp. Data Files 2).

**Figure 3.**
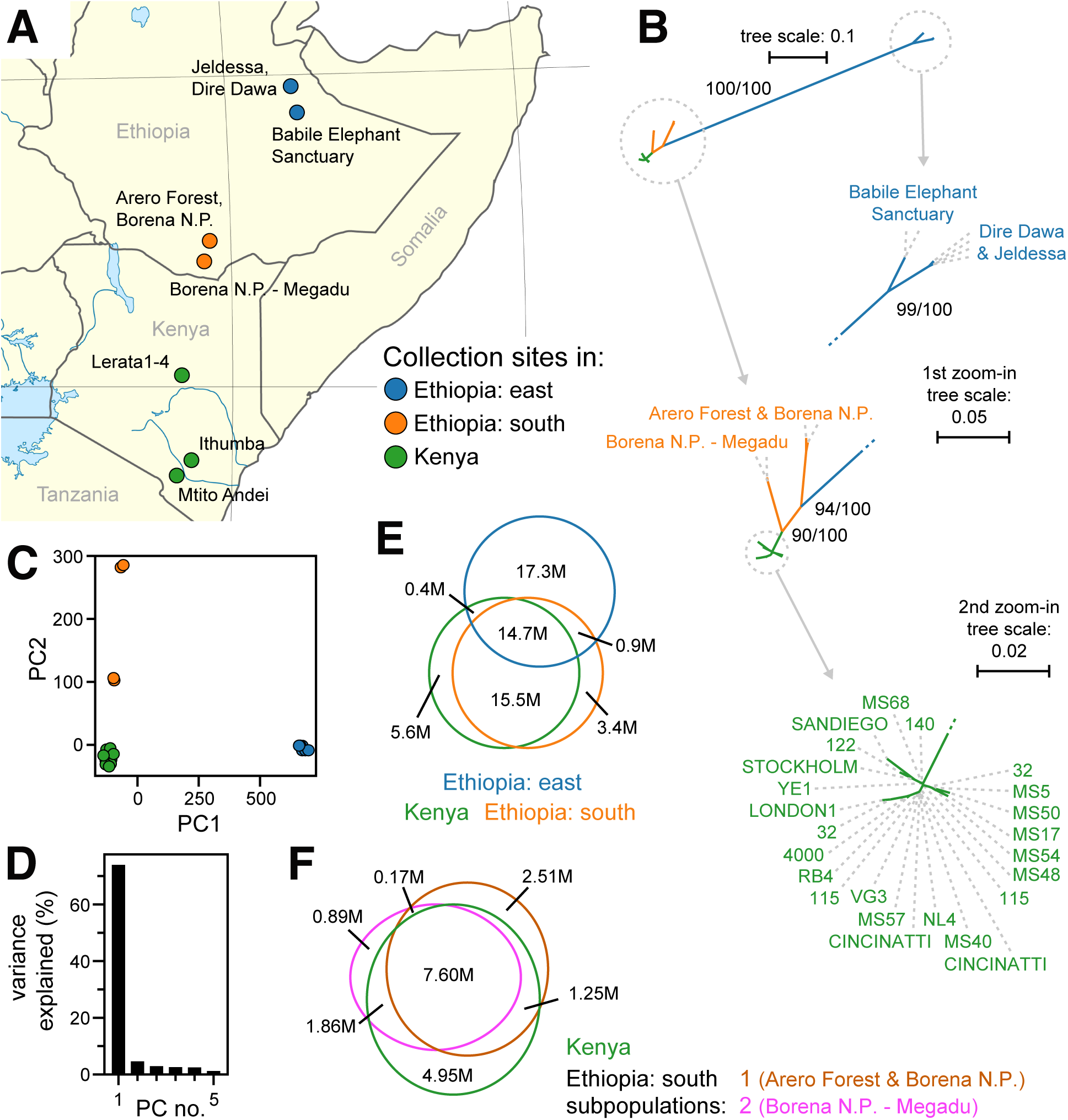
A. A map of collection sites in Kenya and Ethiopia. The captive-bred Kenyan mole-rats were derived from animals collected at the indicated sites (green) in 1979 (Jarvis 1985). The wild-caught Ethiopian samples were collected at the orange (south Ethiopia) and blue (eastern Ethiopia) sites during multiple expeditions from 2008 to 2019 (Zemlemerova et al. 2020). B. Maximum likelihood, unrooted phylogenetic tree for the 34 samples selected from whole-genome sequencing. Nodes with bootstrap support of at least 90% are highlighted. Animals are labelled by their colony name or geographic origin (see Supplemental Table 4) and colored by geographic region as in panel A. C. Principal component analysis of genotype values for all 41 samples. The scatterplot shows the first two principal components (PC1 and PC2) plotted on the x and y axes, respectively. Color coded by geographic region as labelled in panel A. D. Histogram of the variance explained for the first eight principal components (PC1 and PC2 are plotted in panel C). E. Venn diagram of alleles shared across the three geographic regions of sample collection, colored as in panel A. For each polymorphic site, reference and alternative alleles are both represented. Only alleles that were polymorphic across the 34 animals from this analysis are included. F. Venn diagram of allele distributions across the Kenyan and two south Ethiopian populations (Arero Forest and Borena N.P. in brown; Borena N.P. - Megadu in pink; Kenya in green). Only alleles that were polymorphic across the 28 animals from this analysis are included.

To probe population structure, we constructed a maximum likelihood phylogenetic tree (see Methods), including the 24 Kenya-derived animals after kinship-based filtering plus the ten Ethiopian animals. There was high bootstrap support for the Kenyan population being closely related to southern Ethiopian animals, and this combined clade being highly diverged from the eastern Ethiopian animals (Figure 3B). Divergence of the eastern Ethiopian group was also revealed by principal component analysis of genotype variation (Figure 3C). Eastern Ethiopian animals clustered separately from all other animals along the first principal axis, which accounted for >70% of genotype variance (Figure 3C,D). This divergence was also revealed by the distribution of alleles across geographic regions: while the Kenyan and southern Ethiopian animals shared most alleles, the majority of alleles discovered from the eastern Ethiopian animals were unique to that group (Figure 3E; Supplemental Table 7). We quantified the divergence between geography-delimited groups (Kenya, south Ethiopia, east Ethiopia) using the *F_ST_* statistic, which ranges from 0.0 for a free exchange of alleles to 1.0 for no exchange of alleles (Wright, 1943). The eastern Ethiopian group had high *F_ST_* versus either the Kenya-derived group (0.89) or the southern Ethiopian group (0.91), while the southern Ethiopia and Kenya groups shared lower *F_ST_* lower (0.68; Table 1A).

**Table 1.** Population differentiation measured with *F_ST_* statistic, for trios of populations. Values were calculated for bins of 50 adjacent variants, and the genome-wide means and standard deviations are presented. X-linked scaffolds were excluded from this analysis. Sub-table A presents statistics for the Kenya-derived, southern Ethiopian, and eastern Ethiopian populations; sub-table B presents statistics for the Kenya-derived population and the two subpopulations from southern Ethiopia; sub-table C presents statistics for the Kenya-derived population and the two artificially created subpopulations of southern Ethiopian animals (see Methods).

| populations compared | No. sites | $F_{ST}$ mean | $F_{ST}$ SD |
| --- | --- | --- | --- |
| **Sub-table A** |  |  |  |
| three populations | 29688738 | 0.8776 | 0.2149 |
| Kenya vs. south Ethiopia | 12013314 | 0.6780 | 0.3130 |
| Kenya vs. east Ethiopia | 26181861 | 0.8863 | 0.2180 |
| south Ethiopia vs. east Ethiopia | 23918670 | 0.9073 | 0.1582 |
| **Sub-table B** |  |  |  |
| three populations | 29687481 | 0.2670 | 0.3867 |
| Kenya vs. south Ethiopia 1 | 10878434 | 0.6150 | 0.3414 |
| Kenya vs. south Ethiopia 2 | 8978365 | 0.5381 | 0.3286 |
| south Ethiopia 1 vs. 2 | 5820849 | 0.8136 | 0.2079 |
| **Sub-table C** |  |  |  |
| three populations | 29703031 | 0.2474 | 0.3686 |
| Kenya vs. E.south artificial 1 | 11271504 | 0.5958 | 0.3501 |
| Kenya vs. E.south artificial 2 | 11158621 | 0.5930 | 0.3500 |
| E.south artificial 1 vs. 2 | 5821762 | 0.2580 | 0.2808 |

The possibility of admixture between these three population groups was evaluated using the *f_3_*statistic (Patterson et al. 2012), calculated for each of these three geographically-delimited populations with respect to the other two. In all cases, *f_3_* was significantly greater than zero (Table 2A). This result indicated no recent admixture between these groups, with the east Ethiopian population again standing out as the most highly diverged.

**Table 2.** Admixture tests using the *f_3_* statistic. For each trio of populations, the estimate and standard error of the *f_3_* statistic is provided for each population relative to the other two. Sub-table A presents statistics for the Kenya-derived, southern Ethiopian, and eastern Ethiopian populations; sub-table B presents statistics for the Kenya-derived population and the two subpopulations from southern Ethiopia.

| population A | populations B,C | $f_3$ | SE | z-score |
| --- | --- | --- | --- | --- |
| **Sub-table A** |  |  |  |  |
| Ethiopia east | Ethiopia south, Kenya | 0.4991 | 0.00076 | 651.8 |
| Ethiopia south | Ethiopia east, Kenya | 0.01085 | 0.00011 | 103.1 |
| Kenya | Ethiopia south, east | 0.02788 | 0.00013 | 218.9 |
| **Sub-table B** |  |  |  |  |
| Ethiopia south 1 | Ethiopia south 2, Kenya | 0.06355 | 0.00040 | 157.0 |
| Ethiopia south 2 | Ethiopia south 1, Kenya | 0.03617 | 0.00022 | 164.8 |
| Kenya | Ethiopia south 1, 2 | 0.01739 | 0.00016 | 112.2 |

### Allelic distributions suggest two geographically proximal populations in southern Ethiopia to be more closely related to the Kenyan population than to each other

Principal component analysis revealed animals collected from two locales of southern Ethiopia (Borena N.P. and Arero Forest, versus the Megadu block of Borena N.P.; Figure 3A) to be clustered separately from each other, and from Kenya-derived samples, along the second principal axis (Figure 3C). Likewise, phylogenetic clustering placed members of these two populations more proximal to Kenya-derived animals than to one another (Figure 3B). Heterozygosity analysis across the three geographically-defined populations (Kenya, southern Ethiopia, eastern Ethiopia) also supported independent evaluation of the two southern Ethiopian populations. Using high-confidence polymorphisms, the observed versus expected number of heterozygous positions (*H_o_* versus *H_e_*, respectively) were calculated genome-wide, excluding X-linked scaffolds (Table 3A). While observation met expectation for the captive-interbred Kenyan samples (0.29 and 0.27 for *H_o_* and *H_e_*, respectively), substantially less heterozygosity was observed than expected across the southern Ethiopian samples (0.13 versus 0.41 for *H_o_*and *H_e_*, respectively).

**Table 3.** Genetic diversity measured with observed and expected heterozygosity (*H*_o_ and *H*_e_, respectively). Values were calculated for bins of 50 adjacent variants, and the genome-wide mean and standard deviations are presented. X-linked scaffolds were excluded from this analysis. Sub-table A presents statistics for the Kenya-derived, southern Ethiopian, and eastern Ethiopian populations; sub-table B presents statistics for the Kenya-derived population and the two subpopulations from southern Ethiopia; sub-table C presents statistics for the Kenya-derived population and the two artificially created subpopulations of southern Ethiopian animals (see Methods).

|  |  |  | heterzygosity<br>expected |  | heterzygosity<br>observed |  |
| --- | --- | --- | --- | --- | --- | --- |
| pop | No. haploid<br>genomes | No. variants | mean | SD | mean | SD |
| **Sub-table A** |  |  |  |  |  |  |
| **Ethiopia central** | 12 | 4676574 | 0.3397 | 0.0590 | 0.1661 | 0.1380 |
| **Ethiopia south** | 8 | 5821894 | 0.4084 | 0.0560 | 0.1349 | 0.1130 |
| **Kenya** | 48 | 7620360 | 0.2748 | 0.0650 | 0.2875 | 0.0860 |
| **Sub-table B** |  |  |  |  |  |  |
| **Ethiopia south 1** | 4 | 2340806 | 0.4171 | 0.0451 | 0.4229 | 0.2306 |
| **Ethiopia south 2** | 4 | 1182694 | 0.4176 | 0.0499 | 0.4760 | 0.2590 |
| **Kenya** | 48 | 7620360 | 0.2744 | 0.0644 | 0.2866 | 0.0849 |
| **Sub-table C** |  |  |  |  |  |  |
| **E.south artificial 1** | 4 | 94666 | 0.4609 | 0.0383 | 0.1616 | 0.1613 |
| **E.south artificial 2** | 4 | 91505 | 0.4617 | 0.0406 | 0.1589 | 0.1712 |
| **Kenya** | 48 | 148725 | 0.2744 | 0.0644 | 0.2866 | 0.0890 |

We performed additional analyses probing the relationships between the two distinct populations from southern Ethiopia and the Kenya-derived population. The two southern Ethiopian populations were composed of animals collected either from the Arero Forest and northern Borena National Park (s.Ethiopia population 1) or the Megadu block of Borena National Park (s.Ethiopia population 2). The collection sites for these two populations are separated by mountainous terrain and a lava field (Zemlemerova et al., 2021). The *f_3_* statistics (Patterson et al. 2012) were significantly greater than zero for each population versus the other two (Table 2B), indicating no recent admixture. When considered separately, both southern Ethiopian populations had observed heterozygosity that matched expectation (Table 3B). To control for the limited population sizes of these populations (2 animals/4 haplotypes per population), we swapped membership to create two artificial, mis-matched populations. Like the combined southern Ethiopian animal group (Table 3A), these two artificial populations both exhibited far less heterozygosity than would be expected by chance (Table 3C).

Calculation of the *F_ST_* statistic between the Kenya-derived and two southern Ethiopian populations produced somewhat lower values (0.6150 and 0.5381 for populations 1 and 2, respectively; Table 1B) than when those populations were combined (0.6780; Table 1A). Surprisingly, the *F_ST_* statistic between the two southern Ethiopian populations was stantially higher (0.8136; Table 1B), implying a greater barrier to allelic exchange between them versus with Kenyan naked mole-rats. As a control, the swapped/artificial versions of these populations maintained similar values versus the Kenyan population (0.5958 and 0.5930) while drastically lowering the value between populations (0.2580; Table 1C). The notion of the two southern Ethiopian populations being more closely related to Kenyan mole-rat populations than to one another was reinforced by the distribution of polymorphic alleles discovered across these populations: while millions of polymorphisms were discovered for each population that were either unique to it or uniquely shared with the Kenyan population, very few alleles were shared between the two southern Ethiopian populations but not with the Kenyan population (Figure 3F; Supplemental Table 7).

For the eastern Ethiopian samples, phylogenetic clustering (Figure 3B) and observed versus expected heterozygosity (0.16 versus 0.34 for *H_o_* and *H_e_*, respectively; Table 1A) indicated the two geographical populations, deriving from either Jaldessa/Dire Dawa or the Babile Elephant Sanctuary (Figure 3A), to likely also be genetically distinct. However, due to their relative proximity to one another versus their distant relationships to the other populations we studied, this population structure could not have confounded our analysis of the southern Ethiopian and Kenyan populations and so was not analyzed further.

## Discussion

The foundation for our analysis of *H.glaber* genetic diversity was a chromosome-scale assembly of the naked-mole rat genome. We provide this genome assembly, along with the data used to construct it, as resources that may be synergistic with other recent advancements in the quality of *H.glaber* reference genomes (Romanenko et al, 2023; Sokolowski et al, 2024). We additionally provide resources for genome analysis, including *de novo* gene annotations for *H.glaber*, along with transcriptome data supporting them, as well as novel *de novo* genome assemblies for the related species *F.damarensis* and *C.porcellus*.

We measured genetic diversity within a captive-bred *H.glaber* population, as well as across naturally-isolated individuals from either southern or eastern Ethiopia. After their initial collection over 40 years ago and multiple generations of captive breeding, the Kenya-derived collection has maintained substantial genetic diversity. These specimens were relatively closely related to those collected from southern Ethiopia, for which polymorphisms versus our Kenyan reference were identified at a frequency similar to human polymorphisms (∼every 700 basepairs; Supplemental Table 5; Kruglyak & Nickerson, 2001).

In contrast, the eastern Ethiopian specimens were highly diverged from those originating from either Kenya or southern Ethiopia. These results were consistent with previous analyses using a more limited set of genetic markers (Ingram et al. 2015; Zemlemerova et al. 2021). They were also consistent with broad geographic analyses of double digest restriction-site associated DNA markers from the nuclear genome, which also reveals the population group encompassing both Kenya and southern Ethiopia to be highly diverged from the eastern and northern Ethiopian populations (Šumbera et al, 2024). We do not speculate as to whether the degrees of divergence observed might justify novel taxonomic designations within the established *H.glaber* species.

Within southern Ethiopia, substantial division between populations from within versus north of the Megadu block of Borena National Park was strongly supported by our analyses of genome-wide polymorphisms. This distinction may be surprising given the short distance separating collection sites (as little as ∼90km), but it is previously established using a more limited set of genetic sequence and is well justified by the geological barriers separating the populations, including a mountain range and lava field (Zemlemerova et al. 2021).

More surprising was the greater relatedness of each of these southern Ethiopian populations to the captive Kenyan population. Prior analyses suggest the Megadu and Kenyan populations to share a sisterhood relationship versus the Bornea/Arero population (Zemlemerova et al. 2021), which was confirmed in our phylogenetic analysis (Figure 3B). This greater distance was also evidenced by the much larger set of alleles that were unique to the Bornea/Arero population (2.51M) versus the Megadu population (0.89M; Figure 3F). Comparatively few polymorphisms were discovered that were shared between the southern Ethiopian populations but not with the Kenyan population (0.17M; Figure 3F). The captive Kenyan population descends from animals originally collected in central and southern Kenya (Jarvis, 1985), at sites no closer than ∼340km from the nearest Ethiopian collection sites. While the southern Ethiopian populations are separated from one another by natural barriers along the north/south axis, the extended Somali Acacia–Commiphora bushlands to the east provide a potential historical divergence point for those two populations plus the wider Kenyan population. This unexpected distribution of shared genetic diversity may provide insight into the ancient migration patterns that give rise to these populations.

## Methods

### Culture of fibroblast cell-lines

Genomic DNA for the *H.glaber* reference genome was collected from a fibroblast cell line originally derived from skin sections collected from the “stockholm” individual (Hg-01083) as a 1-year-old, non-breeding female. Prior to collection, the skin sections were washed with 70% ethanol. For each skin section, we removed all subcutaneous fat, performed two washes in sterile, cold PBS and performed two washes with cold 70% ethanol. Skin tissue was then cut into small, pea-sized sections, minced with scissors and placed in one well of a sterile, 12-well plate collagenase/Minimum Essential Medium 1x, +Earle’s Salts, +L-glutamine [MEM; Gibco, 11095-080, 500mL] mixture, consisting of 200μL of reconstituted Roche liberase DH research grade purified enzyme blend (Roche, Ref#05401054001) mixed with 1mL of MEM, supplemented with 10% fetal bovine serum [FBS;] and 1x antibiotic/antimycotic [aa; Gibco, 15240-062, 100mL/100x]. Cells were cultured with the same lot number of FBS for the duration of this study. The minced skin and media were mixed with a 5mL serological pipette and then incubated for 3 hours in a 37°C incubator (5% CO_2_, 3% O_2_) with periodical manual agitation to accelerate tissue digestion. After the 3-hour incubation period, the tissue/MEM/collagenase mixture was mixed with a 5mL serological pipette and placed in a 15mL conical tube and centrifuged for 10 minutes at 1000 rpm. The supernatant was then carefully removed, and the remaining pellet was reconstituted in 3mL of 10%FBS/1xaa/MEM media and plated in 1 well of a sterile, 6-well collage-coated tissue culture dish. Plates were incubated in a 32°C incubator (5% CO_2_, 3% O_2_) until confluency. All cells were then trypsinized with 0.25% trypsin-EDTA 1x and passed onto a sterile 10cm collagen-coated tissue culture dish. Skin fibroblasts were cultured until confluence and then split 1:3 on new, sterile collagen-coated tissue culture dishes.

The procedure described above was used to generate the additional *H.glaber* cell lines that were used for genome sequencing (see Supplemental Table 5), as well as the Damaraland mole-rat cell line used for that purpose. The guinea pig dermal fibroblast cell line was purchased from Harlan Sprague, Inc. (Chicago, Ill.). Additional *H.glaber* DNA samples were extracted either similarly-prepared cell lines, flash-frozen lung tissue, or ethanol-stored liver samples (specified below). In all cases, DNA was extracted using a Quick-DNA miniprep kit (Zymo Research, Cat# D3025).

### Whole-genome sequencing

Linked-read sequencing was performed following the 10x linked-reads protocol to prepare Chromium Genome v2 libraries, following the Chromium™ Genome Reagent Kits v2 User Guide (provided at https://support.10xgenomics.com/genome-exome/index). The libraries were sequenced using paired-end (2 x 150bp) sequencing on an Illumina HiSeq 4000. For sorted chromosome sequencing (HiTCH-seq), samples and libraries were prepared from stockholm-derived fibroblasts as described by (Soifer et al. 2020). BioNano optical mapping was also performed from those cells, as described by (Soifer et al. 2020) but only using the DLE-1 Enzyme assay (Bionano Genomics Document 30206A). PacBio long-read sequencing was performed on stockholm fibroblast-derived DNA, using both the RSII and Sequel platforms, and in the latter case using v1.2 and v2.0 chemistries. RSII-sequenced PacBio samples were prepared as described by (Soifer et al. 2020). Sequel-sequenced samples were similarly prepared, with movie time increased up to 10 hours.

### Transcriptome sequencing library prep

For RNA sequencing, total RNA was extracted separately from a variety of tissues, dissected from two animals (a female breeder and a male breeder), as described by (Hilton et al, 2019). Those tissues/organs included: liver, kidney, heart, lung, spleen, pancreas, leg muscle, cerebellum, and whole brain. Equal-mass quantities of RNA from each tissue were pooled for sequencing. For size-selected RNA-seq, total RNA was size selected on a 1% MOPS gel, run for 17 hours, that was then cut into 50 ∼1.5mm slices. RNA from each slice was sequenced on an Illumina HiSeq 4000, as described by (Hilton et al, 2019). Accession numbers for sequence data from each slice (labelled from 1 adjacent to the well and 50 at the bottom of the gel) are available in Supplemental Table 7. PacBio Isoform Sequencing (IsoSeq) was performed on libraries constructed using the Clontech SMARTer cDNA Synthesis Kit and SageELFTM Size-selection System (PN 100-574-400-01) & Diffusion Loading, following manufacturer’s instructions: https://www.pacb.com/wp-content/uploads/2015/09/Procedure-Checklist-Isoform-Sequencing-Iso-Seq-using-the-Clontech-SMARTer-PCR-cDNA-Synthesis-Kit-and-the-BluePippin-Size-Selection-System.pdf.

### *De novo* genome assemblies of the *F.damarensis* and *C.porcellus* genomes

For the Damaraland mole-rat and guinea pig genomes, genomic DNA collected from fibroblast cell lines was sequenced with 10x Genomics linked short-read data to estimated 46X and 45X genome coverage, respectively, as described above. Fold coverage is based on the picogram (pg) estimates of genome size from (Gallardo et al, 2003) and assuming 978 Mb per pg (Dolezel et al, 2003). The 10X Supernova™ Assembler was used to create *de novo* genome assemblies using default settings and manufacturer’s instructions (https://support.10xgenomics.com/de-novo-assembly/software/pipelines/latest/using/running).

### *De novo* genome assembly of the *H.glaber* genome

For *de novo* assembly of the *H.glaber* genome, we followed the Vertebrate Genome Project (Rhie et al. 2021) assembly protocols for genome assembly, phasing, primary polishing using Falcon-unzip, removing duplicated haplotype using purge_dups, scaffolding with scaff10x and BioNano, and secondary polishing with 10x Genomics’ Freebayes. We deviated from the standard VGP1.6 pipeline in four ways: first, all PacBio reads (Sequel 2.0, Sequel 1.2 and RSII) were used in the assembly pipeline; second, for polishing the primary genome assembly with Falcon-unzip, only Sequel1.2 and RSII reads were used (Sequel 2.0 and RSII could not be combined at this step); third, for polishing the secondary genome assembly polish with Freebayes, 10x Genomics linked short-read data was down-sampled to 60X from the original 290X sequence coverage, in order to conform with the specifications of the Vertebrate Genome Project; fourth, only a single round of 10x scaffolding and 10x polishing was executed. At every stage, genome completeness was assessed using BUSCO (Simão et al, 2015) (gene set: Vertebrata_odb9; Supplemental Table 1).

For the grouping of scaffolds that derive from the same chromosome, low-pass sequencing was performed for 3072 flow-sorted chromosomes (deposited in NCBI SRA under BioSample SAMN41435818) and used to assign the 60 >1Mb scaffolds from our assembly into chromosome groups, using the method of (Soifer et al. 2020). Briefly, enrichment scores were calculated estimating the likelihood that each pair of scaffolds derived from the same chromosome. A correlation matrix of these scores (Supp. Data File 1) was used to cluster scaffolds using k-means, and the “Kneedle” algorithm (Satopää et al, 2011), as implemented by the python kneed module (v0.8.5), was used to determine the number of clusters. Because *H.glaber* has 29 autosomes plus sex chromosomes, and the DNA sampled for assembly was from a female, only cluster counts 30 or above were considered, resulting in 32 clusters (scaffold “groups”). These groups were sorted by the summed lengths of component scaffolds. Thirty groups had lengths >10Mb, with the bottom two groups both being singleton scaffolds near the length threshold for inclusion in this analysis: Super-Scaffold_154 (1.69Mb) and Super-Scaffold_183 (1.46Mb). Those scaffolds were designated as ungrouped, leaving 30 groups (Supplemental Table 2) that we suspect correspond to the 30 chromosomes of *H.glaber* (Deuve, 2006). Synteny of the scaffolds with reference mouse (mm10) and human (hg38) genomes was determined using halSynteny (Krasheninnikova et al. 2020). In Figure 1B, syntenic blocks aligned to the same reference genome and separated by less than 0.25 Mb were visually fused.

### Repeat element detection and masking

Genomic repeats specific to *H.glaber* were identified using RepeatModeler (version=1.011), and those were combined with conserved repeats from the DFam database (Hubley et al., 2016). In order to mask repeats for genome analysis, we applied this composite repeat library to the *H.glaber* genome and the other nine genomes used in the whole genome alignment process in RepeatMasker (version=open-4.0.9; flags -s, -xsmall, -poly) (Smit et al, 2013).

### Gene annotation informed by comparative genomics and RNA-seq data

Pacbio IsoSeq data were processed using software tools downloaded from bioconda. Subreads were converted to circular consensus sequences using ccs (version=6.0.0, commit v6.0.0-2- gf165cc26; flags: min-rq=0.9). We removed 51 and 31 cDNA primers, barcodes using lima (version=2.0.0, commit v2.0.0; flags: --dump-clips, --peek-guess). Poly-A tails and artificial concatemers were removed using isoseq3 (version=isoseq3 3.2.2, commit v3.2.2; flags: -- require-polya). The resulting unpolished, full-length, non-concatemer sequences were merged using dataset (version=0.1.27; flags: --force, --TranscriptSet), then transcripts were clustered using the cluster function in isoseq3. IsoSeq reads were aligned to the genome using minimap2 (version=2.17-r941) and the Cupcake pipeline (https://github.com/Magdoll/cDNA_Cupcake/wiki) used to collapse these into a GFF-format file, using the script from that library: collapse_isoforms_by_sam.py.

Size-selected RNA-seq reads were trimmed using TrimGalore (version=0.6.3) and aligned to our *H.glaber* reference genome using STAR (version=2.5.3a).

Gene annotation was performed using the Comparative Annotation Toolkit (CAT) (https://github.com/ComparativeGenomicsToolkit/Comparative-Annotation-Toolkit; Fiddes et al, 2018). Three strategies were applied: 1) liftover of established annotations from the human genome; 2) liftover of established annotations from the mouse genome; and 3) *ab initio* searches, to identify genes that are not conserved in those species. Our three newly-generated genomes (for *H.glaber*, *F.damarensis*, and *C.porcellus*) were simultaneously annotated using CAT, leveraging both comparative genomics (Cactus alignments, described below) and gene expression (both Illumina short-read and PacBio IsoSeq long-read technologies). CAT projects well-curated annotations (here, from mouse and human) onto less well annotated genomes, then cleans and filters those using AUGUSTUS (Stanke et al., 2006). Additional novel information garnered by combining the projected annotations with predictions produced by Comparative Augustus (König et al. 2016) are incorporated into the annotation pipeline and further supplemented and validated by gene expression data. Size-selected RNA-seq was used to inform annotation of intron/exon boundaries (i.e. intron-only mode). IsoSeq data were used in two ways as part of the annotation pipeline: first, as input for AugustusPB; second, after attempted derivation of gene models using Cupcake (described above). Cupcake annotations were used similarly to NCBI annotations -- providing information to Augustus CGP. IsoSeq transcripts were also directly incorporated into the *H.glaber* genome annotation if they provided a novel isoform or locus.

Whole genome alignment was performed using Cactus (version=1.2.3) according to developer instructions (https://github.com/ComparativeGenomicsToolkit/cactus). Ten mammalian genomes were selected for alignment. In addition to the three assemblies generated here (naked mole-rat *H.glaber*, Damaraland mole-rat *F.damarensis*, and guinea pig *C.porcellus*), the following seven reference genomes were included, listed with their UCSC genome browser and NCBI ID’s: human *Homo sapiens* (hg38; GCF_000001405.38; Schneider et al, 2017), mouse *Mus musculus* (mm10; GCF_000001635.26; Church et al, 2009), rat *Rattus norvegicus* (rn6; GCF_000001895.5; Ramdas et al, 2019), rabbit *Oryctolagus cuniculus* (oryCun2; GCA_000003625.1; Carneiro et al, 2014), squirrel *Ictidomys tridecemlineatus* (speTri2; GCA_000236235.1), dog *Canis lupus familiaris* (canFam3; GCA_000002285.2; Lindblad-Toh, 2005), Brandt’s bat *Myotis brandtii* (GCA_000412655.1), and African elephant *Loxodonta africana* (loxAfr3; GCA_000001905.1; Palkopoulou et al, 2018). These species were selected based on their taxonomic relationships to the naked mole-rat and/or the maturity and quality of their genome assemblies and annotations. The Netwick-format string for the alignment-based tree for these genomes is provided in Supplemental Text File 1. Whole-genome alignments are provided as Supp. Data Files 3.

Gene annotations for H.glaber are provided as Supp. Data File 4.

### Karyotype-based estimation of *H.glaber* chromosome sizes

The total size of the *H.glaber* genome was estimated as described in the Results, based on the picogram genome size measured by (Gallardo et al, 2003). Relative sizes of the full chromosomes were calculated from the karyotype image published by Deuve et al (2006) using KICS (Ludwig et al, 2022), with “threshold” set to 0.063 and “blur” set to 0.40.

### Pedigree construction for Kenyan-derived *H.glaber* collection

Our captive *H.glaber* collection descends from 200 animals gathered from six sites in Kenya in 1979 (Figure 3A; Jarvis, 1985) and has since been maintained and interbred in captivity. Throughout the history of the collection, new colonies have been established through the pairing of a single non-breeding male and single non-breeding female by removing them from their birth colonies’ cages and placing them into a new cage. Pairings were considered successful, and a new colony designated, if the female became pregnant. If the male died and there were no surviving/remaining offspring, then the female would sometimes be paired with a new breeding male, under the same colony designation.

As a tool for selection of DNA samples to maximize genetic diversity, we used archival breeding records to construct a pedigree of our Kenya-derived *H.glaber* collection. Supplemental Figure 2 depicts the relatedness between colonies in the *H.glaber* collection as of January, 2020; only network paths connecting the colonies of genotyped individuals and Kenyan founder colonies are shown in Figure 2A. Given the eusocial breeding structure of *H.glaber* society, nodes in this pedigree represent family-colonies (composed of a breeding pair and their offspring). Successful pairing events are shown as arrows to each newly formed colony. The records used to construct this pedigree were incomplete and largely restricted to the prior 25 years, and in only a few instances were we able to trace relationships back to wild-caught progenitors (Figure 2A).

### Genome sequencing and variant calling for *H.glaber* samples

Our reference genome for *H.glaber* was constructed from DNA collected from a single individual from our Kenya-derived collection (“stockholm” a.k.a. Hg-01083; see above). The genomes of 30 additional animals from this collection were sequenced in order to catalog genetic variants (see Supplemental Table 4 for animal data). For each of these animals, DNA was extracted from either a cell line or flash-frozen lung tissue (Supplemental Table 5). Genomes were also sequenced for four wild-caught animals from southern Ethiopia and six wild-caught animals from eastern Ethiopia (Figure 3A; Zemlemerova et al., 2021). For each of these Ethiopian animals, DNA was extracted from ethanol-stored liver samples. For each Kenya-derived and Ethiopian sample, DNA was extracted using a Quick-DNA miniprep kit (Zymo Research, Cat# D3025), and a linked-read library was prepared using the 10X Genomics Chromium™ kit. Paired-end sequencing (2 x 150bp) was performed for these libraries on an Illumina HiSeq 4000.

For variant calling, each sample was aligned to our *H.glaber* stockholm reference genome using Long Ranger™ (version=2.2.2), duplicate reads were filtered using Picard (version=2.19.1), and variants called using HaplotypeCaller in GATK (version=4.0.0). Next, we grouped all samples together and jointly called variants using the GenotypeGVCFs tool in GATK. High-confidence variants were identified by phred quality score (minimum of 30), maximum depth of coverage (0.95 quantile of coverage for each sample), and minimum depth of coverage (whichever number is greater between 6 or the 0.05 quantile of coverage for each sample). Additionally, variants had to be bi-allelic and no more than 20% of samples could have missing data. Variant data are provided in Supp. Data Files 2.

### Kinship analysis

For the sequencing of genetic variants from Kenyan *H.glaber* (described above), 27 of the animals were selected based on our pedigree to maximize diversity. Three full-sibling animals from a single colony, MS5, were also sequenced as positive controls for kinship analysis. The genome-wide panel of high confidence polymorphisms segregating among these 31 animals (30 sampled, plus the stockholm reference; n=7,620,360) was used to estimate pairwise kinship values between these 31 animals. Kinship *r^w^* was measured using the equation:

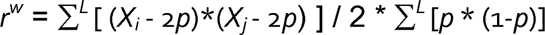

where *X_i_* and *X_j_* are allelic dosages (0,1,2) of individuals *i* and *j*, and *p* is frequency of the non-reference allele in the population at *L* markers (VanRaden 2008; Goudet, Kay, and Weir 2018). All pairwise kinship values are provided in Supplemental Table 6.

For a population in Hardy-Weinberg equilibrium, kinship values are expected to vary between zero and one: zero corresponding to completely unrelated individuals, one corresponding to identical twins. However, violations of Hardy-Weinberg equilibrium, particularly population structure, can also produce negative values. For example, since *p* is calculated across the full population being analyzed, individuals from subpopulations that have distinct *p* values across many alleles, where individual *i* comes from the larger subpopulation (which therefore has greater influence on the overall value of *p*) will be biased towards positive values for the (*X_i_* - 2*p*) term and negative values for the (*X_j_* - 2*p*) term, producing products that are biased in the negative direction.

For Figure 2B, animals were clustered based on kinship using the dissimilarity values of (2 - *r^w^*)^2^, using cluster.hierarchy.linkage (method=’average’) from scipy (v=1.10.1). For any group of samples with kinship estimates >0.5 (the value expected for full siblings), all but one sample was removed for subsequent analyses. There were three such fully-independent groups: group A (animals nmr_1901_5, nmr_1901_20, nmr_1901_21, nmr_1901_14), group B (animals nmr_11, nmr_12, nmr_13), and group C (animals nmr_1901_15, nmr_1901_16). Two additional groups were formed from threshold-exceeding kinship with individual animals from group A: nmr_5 with nmr_1901_20, and nmr_1901_5 with nmr_1909_9. The removal of seven animals (nmr_12, nmr_13, nmr_1901_9, nmr_1901_14, nmr_1901_16, nmr_1901_20, nmr_1901_21) left no kinship values exceeding 0.5, and those 24 remaining animals were used to represent the Kenya-derived population for further analyses.

### Analyses of population divergence, admixture, heterozygosity, and *F_ST_*

Our variant-sequenced *H.glaber* animals included wild-caught animals from eastern Ethiopia (n=6), wild-caught animals from southern Ethiopia (n=4), and captive animals descending from founders caught in Kenya (n=24). Genetic differentiation between these three populations was measured by two methods: 1) construction of a maximum likelihood phylogenetic tree; and 2) principal component (PC) analysis of genotype scores for all samples. A random sample of 1% of the high-confidence polymorphisms (n=299,933) was used to construct a maximum likelihood tree for these 34 *H.glaber* samples using TreeMix v1.13 (Pickrell and Pritchard, 2012), run with the variant block size set to 10. Support for each node in the phylogeny was assessed by running 100 bootstrap replicates and reporting nodes with > 90% support. Branch lengths in Netwick format representation for this tree is provided in Supplemental Text File 1.

For PC analysis, only polymorphisms measured in all samples were used to calculate the first eight PC values using the decomposition function with sklearn python package, version 0.22.2.post1. Evidence of recent admixture was evaluated using the *f_3_* statistic (Patterson et al. 2012), implemented in TreeMix v1.13 (Pickrell and Pritchard 2012), using the ‘threepop’ function with 29.9M polymorphisms, split into blocks of size 10,000.

For Venn diagram construction, for each population being considered, for each polymorphic site for which both alternative alleles were observed across all three populations being considered, the distribution of each alternative allele was determined across the three populations, and the allele was binned according to the combination of populations in which it was observed, at any frequency. Allele counts are therefore twice the number of polymorphic sites contributing to each diagram. For each trio of populations, alleles were only included if their identity was called in all three populations in at least four haplotypes or half of the haplotypes in that population, whichever was greater. Those same criteria were applied when the number of newly-discovered polymorphisms are reported. Raw values for Venn diagrams are provided in Supplemental Table 7.

For each of the three populations, per-site observed heterozygosity *H_o_* was calculated as the number of samples with a heterozygous genotype and the expected heterozygosity *H_e_* as 2*p*(1-*p*) where *p* is the frequency of the non-reference allele in each population. Genetic differentiation between populations *F_ST_* was calculated as the ratio of between-population variance to total variance using an ANOVA. We scored genotypes as binary variables and calculated unbiased estimates of the within- and between-group variance from the one-way ANOVA (Kelly and Hughes, 2019). To summarize these results, the average heterozygosities and *F_ST_* were calculated across non-overlapping windows of 50 polymorphisms, across the genome.

Heterozygosity, *F_ST_* and *f_3_* analyses were performed across geographically defined populations of animals from either Kenya, southern Ethiopia, or eastern Ethiopia, as annotated in Supplemental Table 4, with high-kinship animals excluded from the Kenyan roster as described above. Additional analyses were performed comparing the same Kenyan population to two subpopulations from southern Ethiopia: southern Ethiopian population 1 included animals 3119 and 2953; population 2 included animals 3172 and 3205. As a control against small-number effects, the pairings for the southern Ethiopian subpopulations were swapped, producing two artificial populations. Artificial population 1 included animals 3119 and 3172; artificial population 2 included animals 2953 and 3205.

---

**Supplemental Figure 1.**
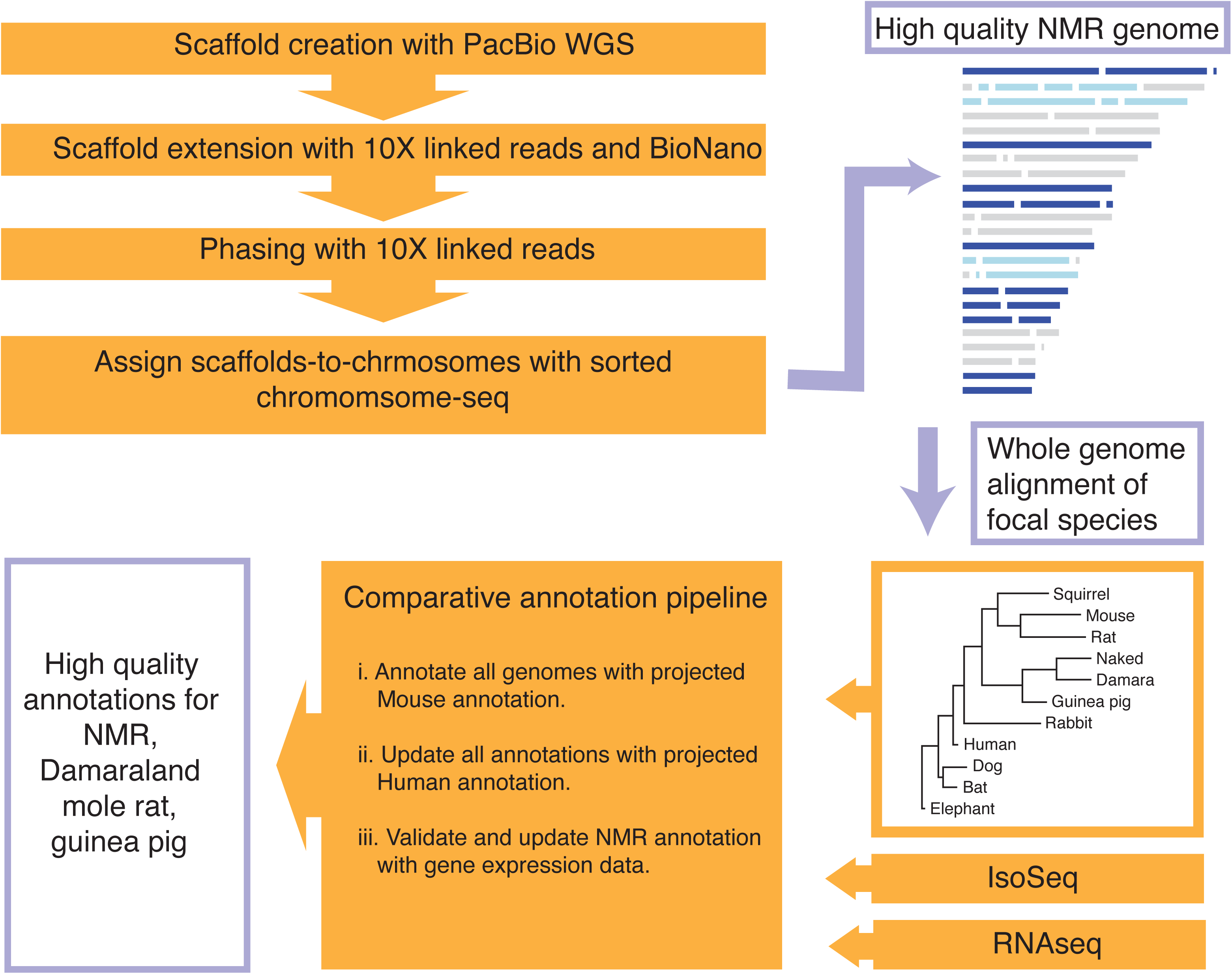
Flow chart describing whole genome assembly and annotation pipelines.

**Supplemental Figure 2.**
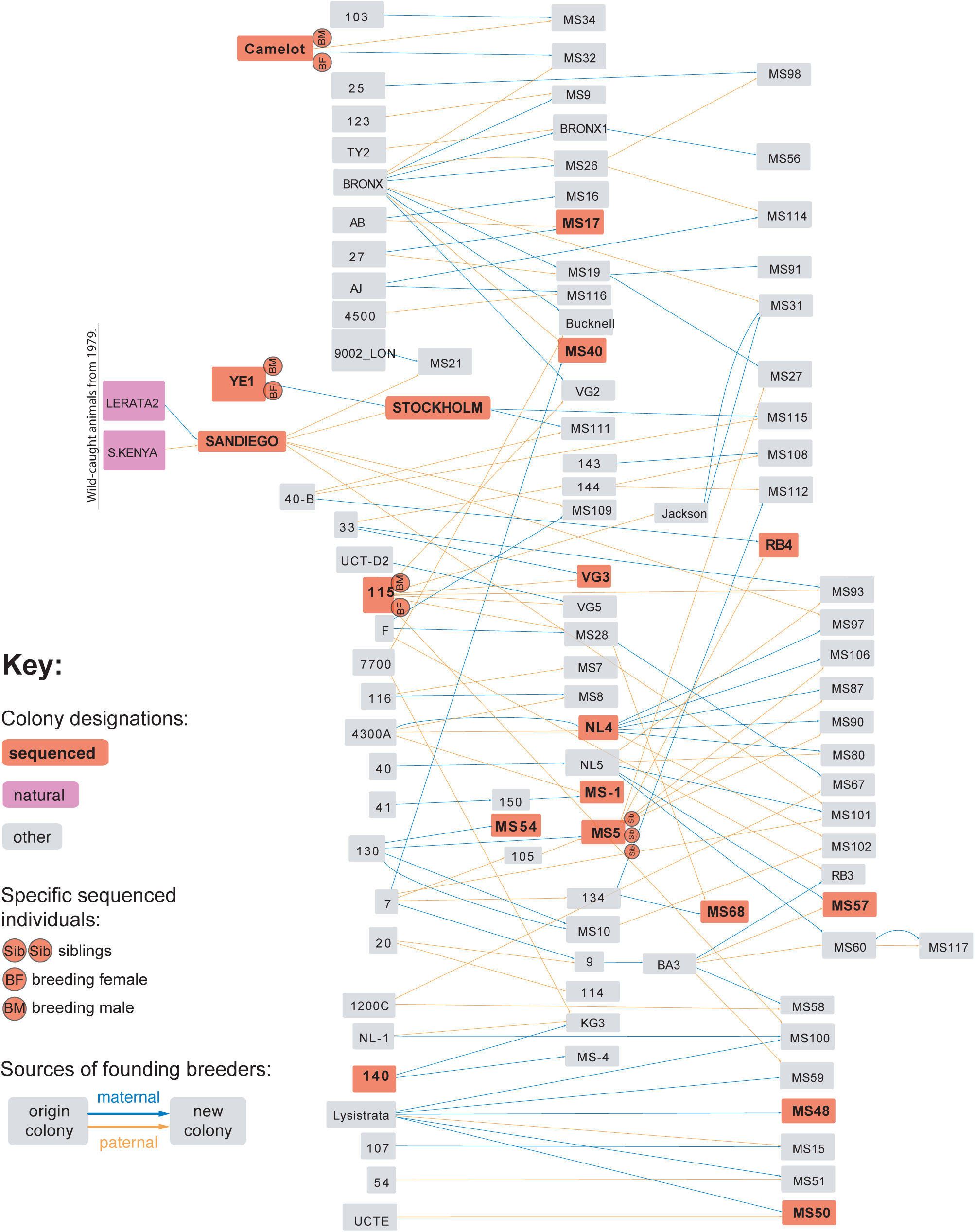
A maximally-complete colony pedigree for our Kenya-derived *H.glaber* collection, constructed using all available historical records. Colored as in Figure 2A, which depicts the subset of this pedigree linking genome-sampled colonies. See Methods for details.

**Supplemental Table 1**

Genome assembly statistics at each assembly step.

**Supplemental Table 2**

Super-scaffold membership in each clustered group, based on chromosome-sorted sequencing (as depicted in Figure 1A).

**Supplemental Table 3**

**Whole genome alignment statistics for 10 focal species. The phylogenetic tree is illustrated in Supplemental Figure 1. Branch lengths in Netwick format for this phylogenetic tree are provided in Supplemental Text File 1.**

**Supplemental Table 4**

Demographic and geographic data for genome-sequenced animals. For ID’s, “Local ID” refers to the animal/sample ID used in this manuscript; where appropriate, “Literature ID” refers to how the animal is referenced in (Ruby et al, 2018 & 2024). For the only Kenya-derived animal not referenced in those publications, a new ID was assigned according to that convention: Hg-08602. Latitude/longitude values refer to the collection sites for Ethiopian specimens: all are north and east, respectively.

**Supplemental Table 5**

Tissue/cell sample type, coverage statistics, and polymorphism counts for each genome-sequenced animal. SRA experiment ID’s are also listed for each animal.

**Supplemental Table 6**

Numerical values for kinship *r^w^*, as displayed in Figure 2B.

**Supplemental Table 7**

Exact values for Venn diagrams from Figure 3.

**Supplemental Table 8**

A table of SRA experiment ID’s for each slice of the size-selected RNA-seq data set (see Methods).

**Supplemental Text File 1**

Netwick format text representations of phylogenetic trees, for genome-aligned species and *H.glaber* individuals from Kenya and Ethiopia.

**Supplemental Data File 1**

A json-formatted pandas data frame of the enrichment scores (estimating the likelihood of each pair of scaffolds deriving from the same chromosome) from flow-sorted chromosome sequencing.

**Supplemental Data Files 2**

Gnu-zipped tar files containing per-animal, per-polymorphism information on either coverage (SuppDataFile2.Depth.tgz; unzips into 63 *.depth.csv files) or genotype (SuppDataFile2.Geno.tgz; unzips into 63 *.geno_score.csv files), for the polymorphisms discovered by whole-genome sequencing. Polymorphisms are organized into files by super-scaffold.

**Supplemental Data Files 3**

Cactus-generated whole-genome alignments of the genome assemblies produced herein for *H.glaber*, *F.damarensis*, and *C.porcellus*, along with published genome assemblies from relevant mammals (see **Methods**). The Gnu-zipped tar files SuppDataFile3.Maf.tgz and SuppDataFile3.Sup.tgz contain the whole-genome alignment (.maf format) and supporting annotation files, respectively.

**Supplemental Data File 4**

A gnu-zipped tar file that contains gene annotation files, in .gff3 and .gp formats.

## Data Resources

The following raw data are available from the SRA under BioProject PRJNA825530: IsoSeq transcriptomics for *H.glaber* (under experiment ID: SRX17869813); size-selected RNA-seq transcriptomics for *H.glaber* (see Supplemental Table 8 for experiment ID’s); short-read sequencing for the reference *H.glaber* genome (stockholm) and Kenya-derived *H.glaber* specimens (see Supplemental Table 5 for experiment ID’s). Additional data for construction of the *H.glaber* reference genome are available from the SRA under BioProject PRJNA1112813, including: HiTCH-seq data (under BioSample SAMN41435818); BioNano optical mapping (under BioSample SAMN41463170); 10X linked reads (under BioSample SAMN41492148); and PacBio long reads (BioSamples SAMN41513184 and SAMN41513185). Data for whole-genome sequencing of the Damaraland mole-rat *F.damarensis* (10X linked reads) are available from the SRA under BioProject PRJNA1110213. Data for whole-genome sequencing of the guinea pig *C.porcellus* (10X linked reads) are available from the SRA under BioProject PRJNA1110269. Data for whole-genome sequencing of Ethiopian *H.glaber* samples for SNP discovery are available from the SRA under BioProject PRJNA1111087.

For *H.glaber*, the Whole Genome Shotgun project has been deposited at DDBJ/ENA/GenBank under the accession JBPUHJ000000000. The version described in this paper is version JBPUHJ010000000.

Supplemental Data Files 1-3, along with other Supplemental Materials, are available from Dryad under the DOI: https://doi.org/10.5061/dryad.m37pvmdf4. The Dryad repository includes raw fasta-formatted versions of the *H.glaber, F.damarensis,* and *C.porcellus* genomes used for analyses presented here.

## Ethics Declarations

Animal experimentation: All of the animals were handled according to approved institutional animal care and use committee (IACUC) protocols, most recently of the Buck Institute (A10138). IACUC approval for housing these colonies of naked mole-rats and Damaraland mole-rats were also obtained from all institutions at which these collections were historically housed.

